# Charge neutralization and β-elimination cleavage mechanism of family 42 L-rhamnose-α-1,4-D-glucuronate lyase revealed using neutron crystallography

**DOI:** 10.1101/2022.08.29.505653

**Authors:** Naomine Yano, Tatsuya Kondo, Katsuhiro Kusaka, Taro Yamada, Takatoshi Arakawa, Tatsuji Sakamoto, Shinya Fushinobu

## Abstract

Gum arabic (GA) is widely used as an emulsion stabilizer and edible coating, and consists of a complex carbohydrate moiety with a rhamnosyl-glucuronate group capping the non-reducing ends. Enzymes that can specifically cleave the glycosidic chains of GA and modify their properties are valuable tools for structural analysis and industrial application. Cryogenic X-ray crystal structure of GA-specific L-rhamnose-α-1,4-D-glucuronate lyase from *Fusarium oxysporum* (FoRham1), belonging to the polysaccharide lyase (PL) family 42, has been previously reported. To determine the specific reaction mechanism based on its hydrogen-containing enzyme structure, we performed joint X-ray/neutron crystallography of FoRham1. Large crystals were grown in the presence of L-rhamnose (a reaction product), and neutron and X-ray diffraction datasets were collected at room temperature up to 1.80 and 1.25 Å resolutions, respectively. The active site contained L-rhamnose and acetate, the latter being a partial analog of glucuronate. Incomplete H/D exchange between Arg166 and acetate suggested that a strong salt-bridge interaction was maintained. Doubly deuteronated His105 and deuteronated Tyr150 supported this interaction. The unusually hydrogen-rich environment functions as a charge neutralizer for glucuronate and stabilizes the oxyanion intermediate. The NE2 atom of His85 was deprotonated and formed a hydrogen bond with the deuterated O1 hydroxy of L-rhamnose, indicating the function of His85 as the base/acid catalyst for bond cleavage via β-elimination. Asp83 functions as a pivot between the two catalytic histidine residues by bridging them, and this His–His–Asp structural motif is conserved in the three PL families.

**Significance Statement:** Although hydrogen transfer plays an important role in enzymatic reactions, hydrogen atoms are generally invisible in macromolecular X-ray crystallography. In the reaction of polysaccharide lyases, substrate activation by negative charge stabilization of uronic acid and base/acid-catalyzed β-elimination reaction have been postulated. Here, we report the neutron crystallography of polysaccharide lyase. Joint X-ray/neutron crystallography of L-rhamnose-α-1,4-D-glucuronate lyase from *Fusarium oxysporum* (FoRham1) complexed with L-rhamnose was performed, and the hydrogen and deuterium atoms were visualized at a high resolution. FoRham1 catalyzes the specific cleavage of the cap structure of gum arabic, which is useful for various applications in the food, cosmetic, and pharmaceutical industries. A detailed catalytic mechanism for FoRham1 was proposed based on the key structural features of its active site.

## Introduction

Arabinogalactan proteins (AGPs) are a family of proteoglycans comprising the plasma membrane, extracellular matrix, and cell walls of various plants (1, 2). AGPs play important roles in many plant physiological processes, including cell death, cell elongation, stress response, intercellular adhesion, and intercellular signal transduction (3). AGPs consist of more than 90% type II arabinogalactan (a carbohydrate moiety) and a core protein rich in hydroxyproline. Type II arabinogalactan contains β-1,3-galactan as the main chain and β-1,6-galactooligosaccharide side chains, which are modified with various sugars, including L-rhamnose (Rha) and D-glucuronic acid (GlcA) (1, 4–6). The complicated polysaccharide structure of AGPs with many branches hampers the detailed analysis of their structure–function relationships (7, 8). Gum arabic (GA) is a subclass of type II arabinogalactan that is sometimes regarded as a representative AGP (6). GA is a sticky exudate from *Acacia* trees that is produced during stress conditions, such as drought and mechanical injury (9). Therefore, GA is widely used as an emulsion stabilizer and coating in various applications, such as food, drink, ink, and pharmaceutical industries (10–12). The primary structure of GA consists of disk-like star-shaped nanoparticles (13). Although the polysaccharide structure of GA has been analyzed using chemical methods and NMR analysis (14, 15), its detailed structure has not yet been elucidated. Because the non-reducing ends of the GA side chains are often capped with α-L-rhamnose-(1→4)-D-glucuronic acid (Rha-GlcA), enzymes that target the disaccharide cap structure would be promising tools for elucidating the detailed structure and function of GA and modifying its physical properties.

We have studied various GA-degrading enzymes from *Fusarium oxysporum* 12S, a phytopathogenic fungus that can grow using GA as the sole carbon source (16–21). L-Rhamnose-α-1,4-D-glucuronate lyase from *F. oxysporum* 12S (FoRham1, EC 4.2.2.-) specifically cleaves the glycosidic bond in the Rha-GlcA moiety of GA and releases Rha (21). FoRham1 generates Δ4,5-unsaturated GlcA via β-elimination. We have previously reported the X-ray crystal structure of FoRham1. The structures of wild-type (WT) enzyme in ligand-free and Rha complex forms were determined at 1.05 and 1.40 Å resolutions, respectively, and the structure of an inactive H105F mutant in complex with Rha-GlcA was determined at 2.42 Å resolution. We identified the active site of FoRham1 on the anterior side of a seven-bladed β-propeller domain and found structural similarities to several polysaccharide lyase (PL) families, including the catalytic residues. Therefore, FoRham1 and its homologs in the former glycoside hydrolase 145 family were reclassified into a novel PL42 family in the Carbohydrate-Active enZYmes database (http://www.cazy.org) (22). Based on structural and mutational analyses, we proposed a possible catalytic mechanism of FoRham1 that involves charge neutralization of uronate carboxyl by Arg166 and proton abstraction and donation by His85 in the lytic cleavage of the O4–C4 bond (21). His105 may support the base-acid dual function of His85. However, this reaction mechanism was proposed for X-ray structures that do not contain hydrogen (H) atoms. Neutron crystallography is a powerful method for determining the H and deuterium (D) atom positions in proteins (23), although it requires a longer beam time and larger volume of crystals (> 1 mm^3^) because the neutron beam intensity is substantially weaker than that of X-rays. Since neutrons are diffracted by atomic nuclei, the amplitude of neutron scattering depends on the structure of the atomic nuclei. H and D atoms contribute to neutron scattering to the same extent as the C, N, O, and S atoms. In this study, we performed joint X-ray/neutron (XN) crystallography of FoRham1 to propose a detailed reaction mechanism based on protonation states around the active site. Although many crystal structures of carbohydrate-active enzymes have been determined using X-ray methods (22), to the best of our knowledge, this is the first report on the neutron structures of more than 40 PL families.

## Results

### Crystallization, diffraction experiments, and joint XN structure determination

The recombinant WT FoRham1 protein expressed in *Pichia pastoris* was used for crystallization. H/D exchange was achieved by changing the solution of purified protein with deuterated buffer, and the crystals were grown using solutions prepared with heavy water. The crystallization conditions were thoroughly re-examined to grow large-volume crystals. A crystal grown to a volume of ∼1.5 mm^3^ in the presence of Rha was used for the multi-probe quantum beam diffraction experiments (Fig. 1*A*; Fig. S1). Time-of-flight (TOF) neutron diffraction data collection (24) was performed using an IBARAKI biological crystal diffractometer (iBIX) in the J-PARC MLF (25–27). The diffraction images were processed using the STARGazer software (28), which employs a profile fitting method for peak integration (29). The same crystal was used for synchrotron X-ray data collection. The resolutions of X-ray and neutron diffraction datasets collected at room temperatures were 1.25 and 1.80 Å, respectively (Table 1). Note that the resolution of the present X-ray data was improved over previously reported data collected at a cryogenic temperature (1.40 Å, PDB ID: 7ESM). The non-hydrogen atoms were modeled according to the X-ray electron density map. Joint XN structure refinement was successfully performed and completed with reasonable statistics for crystallography and protein stereochemistry (Table 1). In the final XN structure, 3130 H atoms, 893 D atoms, and 299 water molecules were modeled (Fig. 1*B*) based on the neutron scattering length density (NSLD) maps. Similar to the previous X-ray structure (21), the C-terminal polyhistidine and c-Myc tags were not modeled due to disorder, whereas a high-mannose *N*-glycan at Asn247 was modeled (Fig. 1*C*). We observed GlcNAc_2_-Man_5_ heptasaccharide in the XN structure (Fig. S2), whereas only tetrasaccharide (GlcNAc_2_-Man_2_) was previously observed in the high-resolution X-ray structure (1.05 Å; PDB ID:7ESK) (21). Two monoatomic cations (Na^+^ and Ca^2+^) were observed at the same positions as those in the previous X-ray structures (21). A Tris molecule, which was derived from the reservoir solution for growing huge crystals, was bound on the surface of the posterior side (Fig. S3*A*). The two cations and Tris were located away from the active site and do not participate in substrate-binding or catalysis. Rha and an acetate ion were bound at subsites −1 and +1, respectively, as observed in the previous X-ray structure of WT complexed with Rha (Fig. 1*D*; Fig. S3*B* and Fig. S3*C*) (21).

**Table 1.**
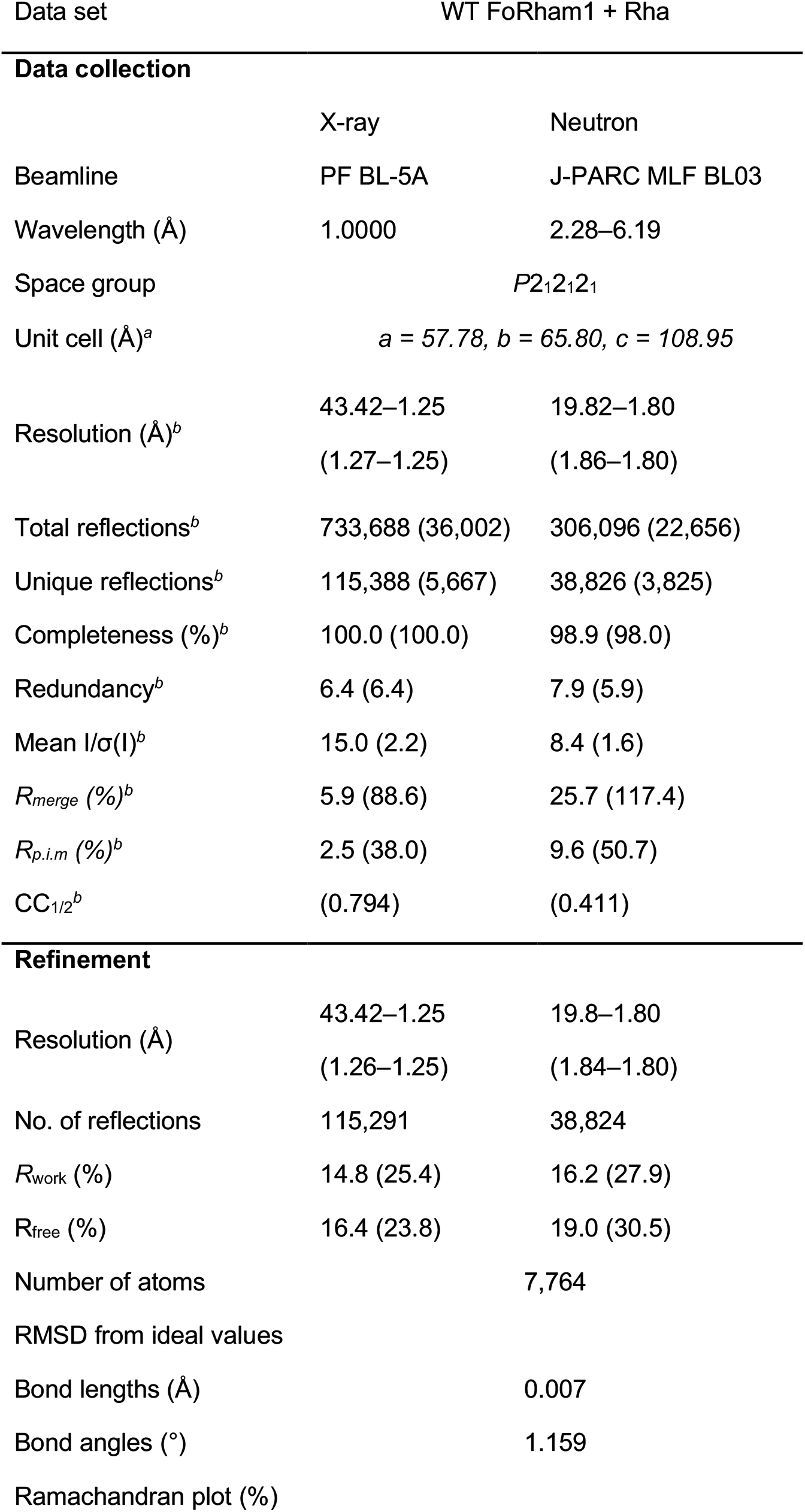
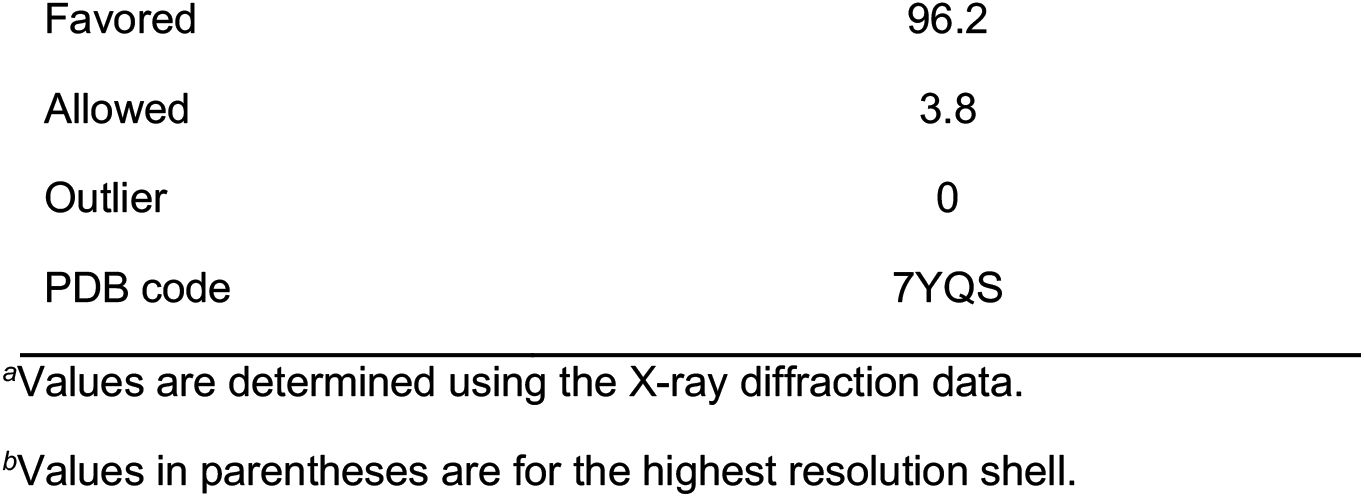
Crystallographic data collection and refinement statistics.

**Data collection**
|  | X-ray | Neutron |
| --- | --- | --- |
| Beamline | PF BL-5A | J-PARC MLF BL03 |
| Wavelength (Å) | 1.0000 | 2.28–6.19 |
| Space group | $P2_12_12_1$ | |
| Unit cell (Å)<sup>a</sup> | $a = 57.78, b = 65.80, c = 108.95$ | |
| Resolution (Å)<sup>b</sup> | 43.42–1.25<br>(1.27–1.25) | 19.82–1.80<br>(1.86–1.80) |
| Total reflections<sup>b</sup> | 733,688 (36,002) | 306,096 (22,656) |
| Unique reflections<sup>b</sup> | 115,388 (5,667) | 38,826 (3,825) |
| Completeness (%)<sup>b</sup> | 100.0 (100.0) | 98.9 (98.0) |
| Redundancy<sup>b</sup> | 6.4 (6.4) | 7.9 (5.9) |
| Mean I/σ(I)<sup>b</sup> | 15.0 (2.2) | 8.4 (1.6) |
| $R_{merge}$ (%)<sup>b</sup> | 5.9 (88.6) | 25.7 (117.4) |
| $R_{p.i.m}$ (%)<sup>b</sup> | 2.5 (38.0) | 9.6 (50.7) |
| CC<sub>1/2</sub><sup>b</sup> | (0.794) | (0.411) |
| **Refinement** |  |  |
| Resolution (Å) | 43.42–1.25<br>(1.26–1.25) | 19.8–1.80<br>(1.84–1.80) |
| No. of reflections | 115,291 | 38,824 |
| $R_{work}$ (%) | 14.8 (25.4) | 16.2 (27.9) |
| $R_{free}$ (%) | 16.4 (23.8) | 19.0 (30.5) |
| Number of atoms | 7,764 |  |
| RMSD from ideal values |  |  |
| Bond lengths (Å) | 0.007 |  |
| Bond angles (°) | 1.159 |  |
| Ramachandran plot (%) |  |  |

|  |  |
| --- | --- |
| Favored | 96.2 |
| Allowed | 3.8 |
| Outlier | 0 |
| PDB code | 7YQS |

<sup>a</sup>Values are determined using the X-ray diffraction data.
<sup>b</sup>Values in parentheses are for the highest resolution shell.

**Fig. 1.**
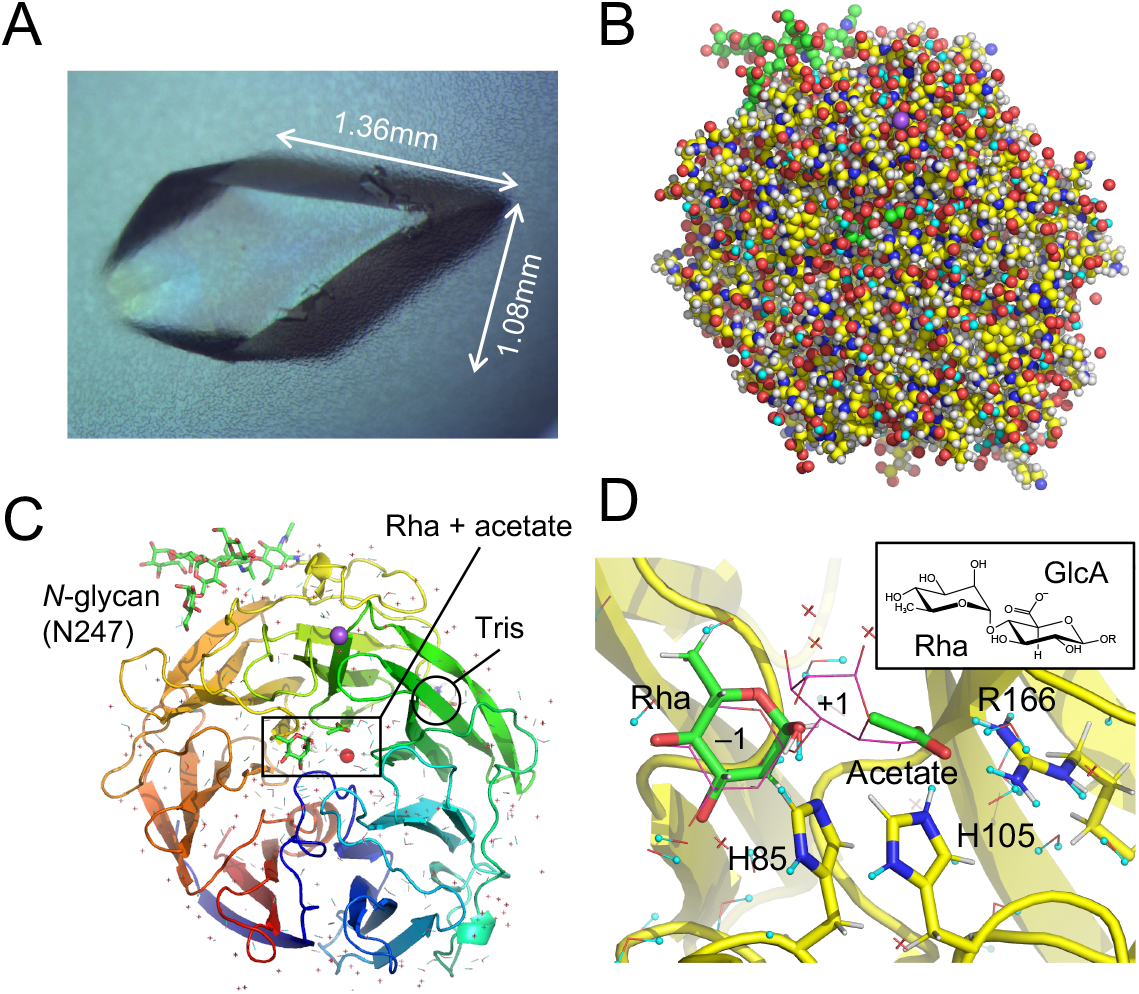
Joint X-ray/neutron (XN) structure of L-rhamnose-α-1,4-D-glucuronate lyase from *Fusarium oxysporum* (FoRham1). (*A*) Huge crystal used in crystallography. (*B*) Overall structure is shown by sphere representation. Atoms are colored as follows: C (yellow for protein and green for ligands and *N*-glycan), O (red), N (blue), Na (purple), H (white), and D (cyan). (*C*) Overall structure is shown by a rainbow-colored ribbon representation viewed from the anterior side of the β-propeller domain. Red sphere and yellow sticks indicate the Ca^2+^ ion and Tris bound on the posterior side, respectively. Water molecules are shown by crosshairs (no H or D atoms modeled) or lines. (*D*) Active site is shown by sticks of the ligand (Rha and acetate at subsite –1 and +1) and the catalytic residues. The substrate (Rha-GlcA) in the X-ray structure (PDB ID: 7ESN) is superimposed (magenta lines). H and D atoms are colored in white and cyan, respectively, and protons are shown as spheres. The chemical structure of Rha-GlcA is shown in the *inset*.

### Rha and acetate in the active site

Fig. 2*A* shows the NSLD map of Rha. The protonation states near the hydroxy groups of Rha were determined by the density peaks that appeared via H/D exchange. O1 atom of Rha is located within the hydrogen bond distance to the NE2 atom of His85 and the OH atom of Tyr150. The D atom involved in the hydrogen bond between O1 of Rha and NE2 of His85 was located at a covalent bond distance with the O1 atom. The D:H occupancy ratio of this atom was 0.60:0.40. A slight positive peak observed between O1 of Rha and wat841 suggested that the O1-D bond of Rha possibly takes a multi-conformer, but the peak height was significantly lower than that near His85. There was no positive peak between O1 of Rha and Tyr150 OH, whereas the OH of Tyr150 formed a hydrogen bond with acetate. XN structure of the Rha complex, which mimics the reaction product, strongly suggests that the proton donor for glycosidic bond cleavage is His85, not Tyr150. At the HE1 atom of His85, a positive peak (D:H = 0.60:0.40) was observed in the NSLD map (Fig. 2*A*, lower panel), suggesting that H/D exchange occurred at the C–H bond. Exchange of the HE1 atom of histidine residues with DE1 has been reported in the neutron crystallography of myoglobin (23, 30) and catalase (31). H/D exchange of the C–H bond was also observed in the guanine base of Z-DNA (32). Arg331 is hydrogen-bonded to O2 and O3 in Rha (Fig. 2*A*, upper panel). Positive peaks near the NH1 and NH2 atoms of Arg331 and O2 of Rha were presented, and the D:H ratios of these atoms were 0.65:0.35, 0.67:0.33, and 0.65:0.35, respectively.

**Fig. 2.**
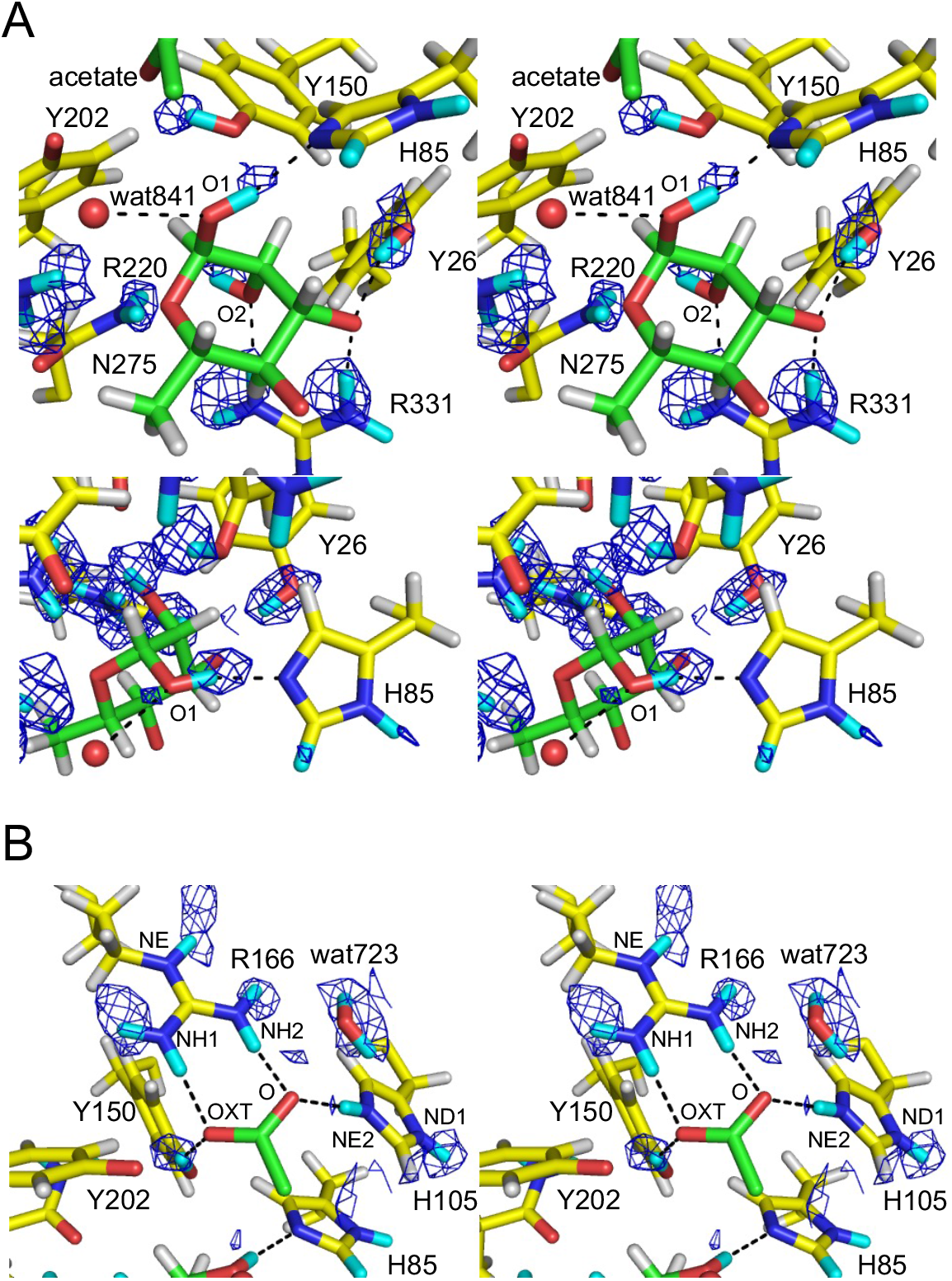
Stereo view of the L-rhamnose (Rha)- and acetate-binding sites with *mF*_o_-*DF*_c_ neutron scattering length density (NSLD) maps (blue mesh). (*A*) Rha-binding site centered at Rha (upper panel, 3.0σ) and His85 (lower panel 2.5σ). Following atoms were excluded from map calculation: DO1 and DO2 of Rha; DH1 of Tyr26; DD1, DE1, and DE2 of His85; DH of Tyr150; DE, DH11, DH12, DH21, and DH22 of Arg220; DD1 and DD2 of Asn275; and DE, DH11, DH12, DH21, and DH22 of Arg331. (*B*) Acetate-binding site with NSLD map (2.9σ). Following atoms were excluded from map calculation: DD1 and DE2 of His105; DH of Tyr150; DE, DH11, DH12, DH21, and DH22 of Arg166; and D1 and D2 atoms of wat52.

The position of acetate corresponds to GlcA carboxylate at subsite +1 (Fig. 1*D*). Acetate is within the hydrogen bond distances to NE2 of His105, OH of Tyr150, NH1 and NH2 of Arg166, and OH of Tyr202 (Fig. 2*B*). A strong positive peak (D:H = 0.65:0.35) was observed near the OH of Tyr150, and this D atom formed hydrogen bonds with the OXT atom of acetate, indicating that Tyr150 was deuteronated (protonated). There was no positive peak between acetate and Tyr202, indicating that there was no hydrogen bond between OH of Tyr202 and OXT of acetate (distance = 2.6 Å). No strong positive peaks were found between the acetate and side chain nitrogen atoms of Arg166 (NH1 and NH2) at 2.9σ level, whereas positive peaks of DE11, DH21, and DE atoms of Arg166 were observed. Yonezawa et al. proposed that Arg52 in photoactive yellow protein, which is hydrogen-bonded with tyrosine and threonine side chains, adopts an electrically neutral form (33). However, we inferred that the Arg166 side chain was positively-charged because of the apparent presence of a salt bridge with acetate. When H/D exchange is incomplete, the negative and positive peak contributions of the H and D atoms cancel each other to yield no or weak peaks in the NSLD maps (23). The calculated D:H occupancy ratios of the atoms between the acetate and the NH1 and NH2 atoms were 0.55:0.45 and 0.50:0.50, respectively, and a very weak positive peak was observed near the NH1 atom at the 2.4σ level (Fig. S4*A*). Because the salt-bridged local structure of Arg166-acetate is similar to that of the Arg–Glu/Asp interactions in proteins, we examined the NSLD map around the salt bridge interaction between Arg112 and Glu76 for comparison (Fig. S4*B*). In the Arg112 side chain, positive peaks were observed near the NH1 and NH2 atoms at 2.8σ level (D:H = 0.74:0.26 and 0.66:0.34 for DH12 and DH22, respectively), indicating that H atoms were efficiently exchanged with D atoms in this area. The D:H occupancy ratios of refined structures via joint XN crystallography can be used as an indicator to interpret the strength of hydrogen bonds and local stability (34). Therefore, the lower D:H occupancy ratio of the atoms between Arg166 and acetate compared to the reference Arg–Glu interaction may indicate the presence of a strong salt bridge interaction, resulting in an incomplete H/D exchange. Between the O atom of acetate and NE2 of His105, a positive peak near His105 (D:H = 0.51:0.49) was observed (Fig. 2*B*). Because a positive peak was also observed near ND1 of His105, this histidine was shown to have a positive charge, as it was doubly deuteronated. The weaker positive peak near NE2 of His105 than ND1 also seems to be due to the strong salt bridge interaction with acetate. The positively charged side chains of Arg166 and His105 in subsite +1 facilitate substrate binding and stabilize the negative charge of GlcA carboxylates.

### Mutational analysis

In our previous report, site-directed mutants of several residues, including His85, Arg166, His105, and Tyr150, in the active site of FoRham1 were analyzed, and these residues were found to be important for enzyme activity (21). In this study, we focused on Tyr202 near the acetate ion (Fig. 2*B*). We noticed that Asp83 may play an important role because it forms hydrogen bonds with the catalytically essential “double histidine” residues, His85 and His105 (Fig. 3*A*). We prepared site-directed mutants of Asp83 and Tyr202. The activities of D83A, D83E, Y202A, Y202F, and Y202W toward GA were not detected, and D83N retained 13% of its activity compared with the WT (Fig. S5*A*). In contrast to the WT, with an optimal pH of 7.5, the activity of D83N increased at a higher pH (Fig. S5*B*). This result suggests that Asp83 holds the two catalytic histidine residues at precise positions and controls their charge states in the deprotonated state, thereby supporting enzyme catalysis. Moreover, the side-chain hydroxy group of Tyr202 was shown to be essential for this activity. As there was no positive peak near the OH atom of Tyr202 (Fig. 2), we presumed that this residue was deprotonated. The positively-charged environment between Arg166 and Arg220 may stabilize the deprotonated (negatively-charged) state of Tyr202 and lower its p*K*_a_ (Fig. 3*A*). Although the detailed function of Tyr202 in enzyme catalysis remains elusive, its deprotonated side chain may maintain a charge balance in the active site that has two arginine residues in proximity.

**Fig. 3.**
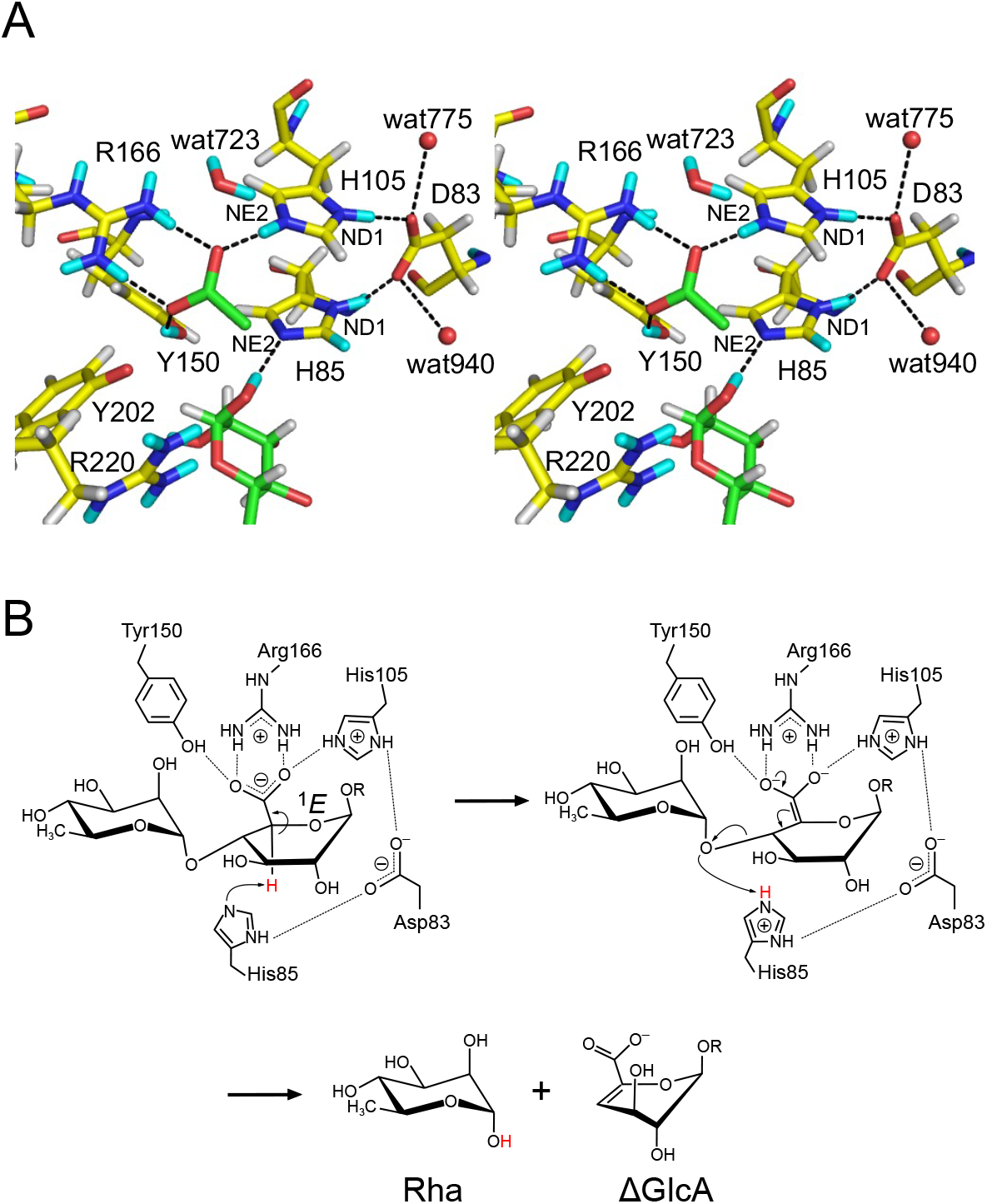
(A) Catalytic residues in the active site and (*B*) proposed reaction mechanism. (*A*) Stereo view of the Rha- and acetate-binding sites including Asp83. D and H atoms (cyan and white) determined via joint XN crystallography are shown. (*B*) Two-step *syn*-elimination mechanism is shown with catalytically important residues. R = H or another sugar residue.

## Discussion

### His–His–Asp triad and possible H/D exchange mechanism

The double histidine motif is present not only in the PL42 family, but also in other PL families (21). Here, we showed that Asp83, which bridges histidine residues behind the active site, is also essential for the enzyme activity. The His–His–Asp triad motif is well-conserved in bacterial ulvan lyases belonging to PL24 and PL25 families (Fig. S6) (35–37). The carboxy group of Asp83 was hydrogen-bonded to the ND1 atoms of His85 and His105 (Fig. 3*A*). The D:H occupancy ratio of these atoms in His85 and His105 were 0.55:0.45 and 0.68:0.32, respectively, indicating that the hydrogen atoms in the two histidine residues were deuterated. In the crystal structure, two water molecules were hydrogen-bonded to Asp83, and a water molecule was found in the vicinity of NE2 of His105. A possible H/D exchange mechanism by solvent molecules is summarized in Fig. S7. The frequent interconversion of the protonation states of His85 with nearby hydrogens (or deuterium) is suitable for its catalytic role as a general base/acid.

### Proposed enzymatic reaction mechanism

The updated catalytic mechanism for FoRham1 proposed in this study is shown in Fig. 3*B*. This mechanism is based on a stepwise *syn*-elimination pathway, which is often proposed for metal-independent PLs (38, 39). The function of Arg166 as a charge neutralizer is enhanced by the side chains of the positively-charged His105 and protonated Tyr150. The unusual environment of the GlcA carboxylate with four hydrogen bond donors (two from Arg166 and one each from His105 and Tyr150) increased the reactivity of the C5 atom and stabilized the oxyanions of an “electron sink” intermediate that is formed after proton abstraction. His85 functions as a catalytic base in the first step because the NE2 atom is deprotonated, as revealed in this study. A distorted envelope conformation (^1^*E*) of the GlcA pyranose ring alleviates the steric hindrance around the H5 atom and brings the hydrogen atom closer to His85 (Fig. 1*D*) (21). In the second step, His85 functions as a general acid to donate a proton to the glycosidic bond oxygen. His85 is connected to His105 and water molecules through hydrogen bonds via Asp83. This hydrogen-bond network possibly modulates the p*K*_a_ of His85 at each catalytic step and supports the dual function of base/acid catalysis.

### Conclusion

Joint XN crystallography of FoRham1 using high-resolution data was used to visualize the positions of the D and H atoms in the crystal complexed with the reaction product, Rha. Fortunately, an acetate ion was bound to subsite +1 as a mimic of the carboxylate group of GlcA to illustrate the unusual hydrogen-rich environment at this site. Direct observation of the D atoms around O1 of Rha excluded Tyr150 from candidates of the catalytic residue in the glycosidic bond cleavage, providing evidence that His85 is the catalytic acid. The deuteronated state of Tyr150 suggests that the catalytic base for the first reaction step is unlikely. This is in contrast to the case of the PL25 ulvan lyase PLSV_3936, where Tyr188 (equivalent to Tyr150) was suggested to function as the catalytic base (37). The “second” histidine, His105, supports the charge neutralizer, Arg166, and modulates the catalytic base/acid functions of His85 through a hydrogen bond network. Asp83 seems to play a pivotal role in controlling the protonation states of the double histidine residues, and the His–His–Asp triad motif is conserved among the PL42, PL24, and PL25 families. The hydrogen bond chain between the histidine residues of FoRham1 is reminiscent of the “Newton’s cradle” proton relay between the catalytic base and acid residues of an anomer-inverting glycoside hydrolase, *Pc*Cel45A (40). Further studies on the visualization of hydrogen/deuterium atoms in numerous carbohydrate-active enzymes will reveal their precise regulatory mechanisms in intricate three-dimensional structures.

## Materials and Methods

### Preparation of huge protein crystals

The recombinant protein expressed in *P. pastoris* with a C-terminal 6× histidine and c-Myc tag was purified as previously described (21). In a previous cryogenic X-ray diffraction experiment, crystals of WT complexed with Rha were obtained via the sitting-drop vapor diffusion method with a 96-well crystallization plate. The crystallization drop was made by mixing the protein solution (24 mg/mL protein, 20 mM Tris-HCl pH 8.0, and 0.1 M Rha) with the reservoir solution of PEG Rx HT #66 (0.1 M Bicine pH 8.5, 10% [v/v] 2-propanol and 30% [w/v] PEG1500) that was purchased from Hampton research. Because the crystals did not grow in homemade crystallization solutions (21), the crystallization conditions were re-examined to obtain large volume crystals for neutron diffraction experiments. To facilitate H/D exchange of the protein and reduce background scattering from H atoms in the diffraction pattern, reagents with heavy water (99.9% D_2_O) were used for crystallization. The purified protein solution was exchanged with 20 mM Tris pD 8.0 buffer using Amicon Ultracel-10K and concentrated to 24 mg/mL. A large-scale sitting-drop vapor diffusion sitting drop method was adopted using a Falcon 60 × 15 mm center well organ culture dish (Corning Inc., Corning, New York) (31). In the outside well of the culture dish, 2 mL of reservoir solution consisting of 0.1 M Tris pD 8.5, 0.1 M Rha, and 33.0% (w/v) PEG1500 in heavy water was poured into the outside well of the culture dish. A stock solution of 1 M Rha, dissolved in heavy water, was used to prepare the reservoir solution. A siliconized glass cover slide was placed in the center well of the organ culture dish. A drop prepared by mixing 100 μL of protein solution and 100 μL of reservoir solution was placed on a cover slide. The culture dish was closed with a lid using high-vacuum grease (Dow Toray Co., Ltd., Tokyo, Japan) as a sealer and incubated at 20 °C for ∼10 weeks until the crystals grew to a size of > 1 mm^3^.

### TOF neutron diffraction experiments

The crystal of WT complexed with Rha was sealed in a quartz capillary (Hilgenberg GmbH, Malsfeld, Germany) with a custom-made stainless-steel magnet base and 20 μL of the reservoir solution (Fig. S1*B*). TOF neutron diffraction experiments were performed using a BL03 iBIX diffractometer at J-PARC MLF (Tokai, Japan) (25–27) with 34 two-dimensional position-sensitive detectors (41) at room temperature. The accelerator power of the proton beam for the spallation neutron source was 600 kW. A neutron diffraction dataset was collected using a circular beam with a diameter of 3 mm at room temperature and a selected neutron wavelength of 2.28–6.19 Å. The capillary sample was placed on a 3-axes goniometer and exposed to 736,000 pulsed neutrons for each crystal orientation. In total, 34 goniometer settings were selected to collect the entire dataset required for structure refinement. Incoherent neutron scattering data from a 4.8-mm vanadium sphere were collected using the same neutron wavelength range as the protein crystal. This procedure was performed to correct for the variance in the detection efficiency of pixels within one detector, the difference in neutron beam intensities by wavelength, and the difference in detection efficiency by wavelength. Data reduction was performed using STARGazer 3.4.3 (28). Data statistics were calculated using the unit cell constants determined via X-ray diffraction.

### ray diffraction experiments

After neutron diffraction data measurements, the same crystal was used for the synchrotron X-ray diffraction experiment on BL-5A at the Photon Factory of the High Energy Accelerator Research Institute (KEK, Tsukuba, Ibaraki, Japan). The crystal was exposed to an X-ray beam of 1.0 Å wavelength with a size of 20 μm × 20 μm at 10% of the maximum intensity at room temperature. Diffraction images of 180 frames were obtained using the oscillation method with 1.0º steps and 1.0-second exposure per frame. The diffraction images were processed using XDS (42). The statistics were calculated using AIMLESS (43).

### Joint XN refinement

Joint XN refinement was performed using PHENIX 1.17.1_3660 (44, 45) and Coot programs (46). Using the reflection tool in PHENIX, the X-ray and neutron diffraction datasets were merged into an MTZ format file. Five percent of the reflections commonly existing in both datasets were randomly assigned as a test dataset for cross-validation. The initial structure model was solved via the molecular replacement method using X-ray data and the atomic coordinates of WT FoRham1 (PDB ID: 7ESK). The solvent water molecules and ligands were then removed from the atomic coordinates. After several cycles of refinement using the X-ray data, the atoms of water, oxygen, Na^+^, Ca^2+^, Tris, Rha, and acetate were placed, and joint XN refinement was performed. The acetate ion was probably derived from metabolites of the *P. pastoris* culture. After several cycles of refinement, H and D atoms were generated in the model using phenix.readyset. The exchangeable hydrogen site atoms were treated as disordered models of H and D atoms, and the initial occupancies were set at 0.5/0.5. Initially, exchangeable site atoms other than those in the main chains were removed from the atomic coordinates. After several cycles of refinement using this model, the H and D atoms at the exchangeable sites of the protein side chains, water, and ligands were added to the atomic coordinates to include them in the refinement. H/D atoms were added when exchangeable hydrogen sites were observed in the NSLD map, or when non-hydrogen atoms bonded to exchangeable hydrogens were clearly observed in the X-ray electron density map. This procedure was repeated several times. The data collection and refinement statistics are summarized in Table 1. All molecular figures were drawn using PyMOL (Schrödinger LLC).

### Site-directed mutant analysis

Site-directed mutagenesis to construct the variants was performed using the PrimeSTAR mutagenesis basal kit (Takara Bio Inc., Kusatsu, Shiga, Japan) with the pPICZαA vector containing the mature *Forham1* gene as the template. The primers used here are shown in Table S1. Recombinant enzymes expressed in *P. pastoris* were purified and assayed, as previously described (21). The assay method is described in Fig. S5.

## Acknowledgments

We would like to thank the staff of the Structural Biology Research Center and Photon Factory at KEK, and Dr. Chihaya Yamada, Mr. Hiromu Arakawa, and Dr. Toma Kashima for X-ray data collection, the staff of iBIX and MLF at J-PARC for neutron data collection, and Dr. Michiko Konno for her support with neutron crystallography. The neutron experiment was conducted as a project for the Ibaraki Prefectural Local Government Beam Line at J-PARC MLF (Proposal Nos. 2019PX3015 and 2020PX3018). The X-ray experiments were conducted with the approval of the Photon Factory Program Advisory Committee (Proposal Nos. 2018G043, 2019G017, 2019G018, 2020G021, and 2019G148). This study was partially supported by JSPS-KAKENHI (19H00929, 15H02443, and 26660083 to S.F.).

