## Supporting Information for "Charge neutralization and β-elimination cleavage mechanism of family 42 L-rhamnose-α-1,4-D-glucuronate lyase revealed using neutron crystallography"

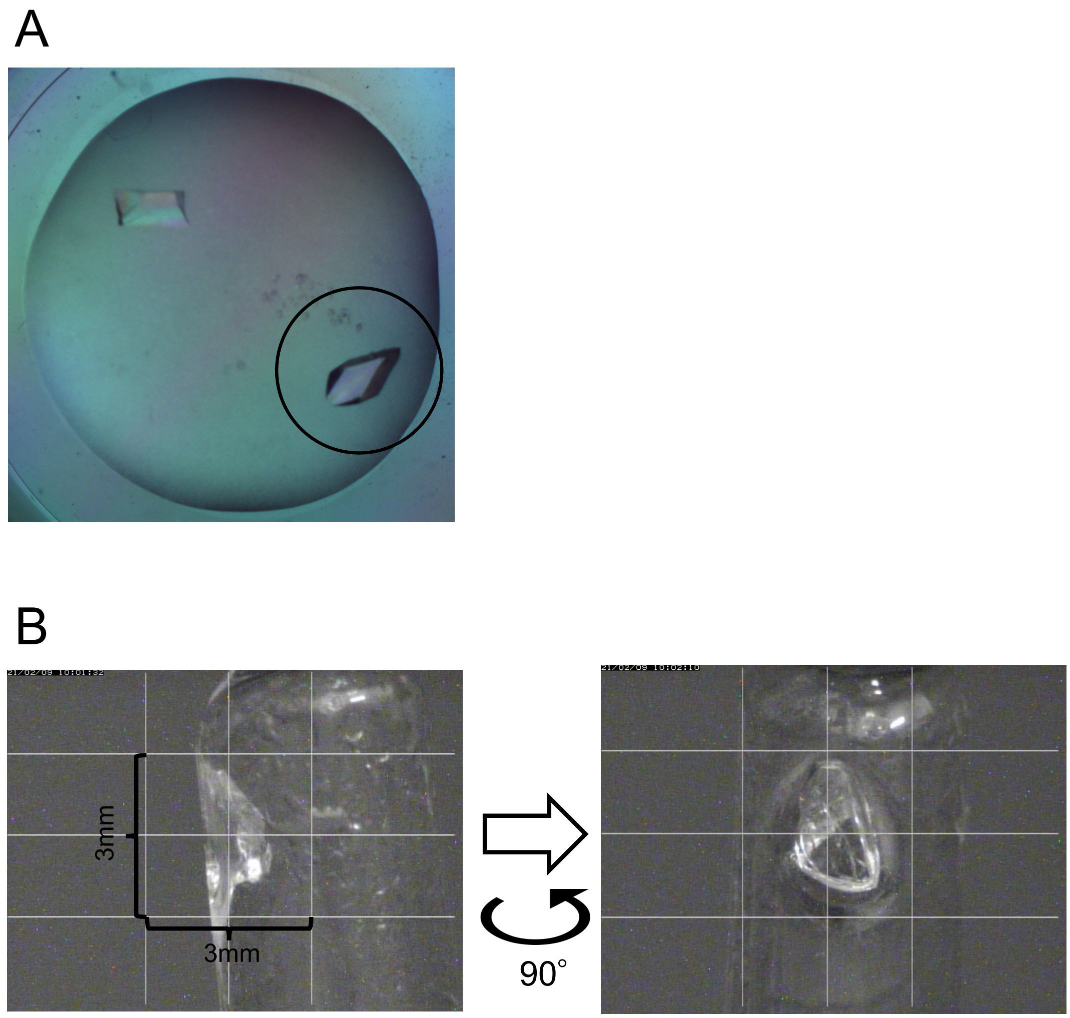


**Fig. S1.** Photographs of the huge crystal. (*A*) The crystallization drop. Initial drop volume was 200 μL. The crystal used for the diffraction experiments (Fig. 1*A*) is circled. (*B*) The crystal was sealed in a quartz capillary.

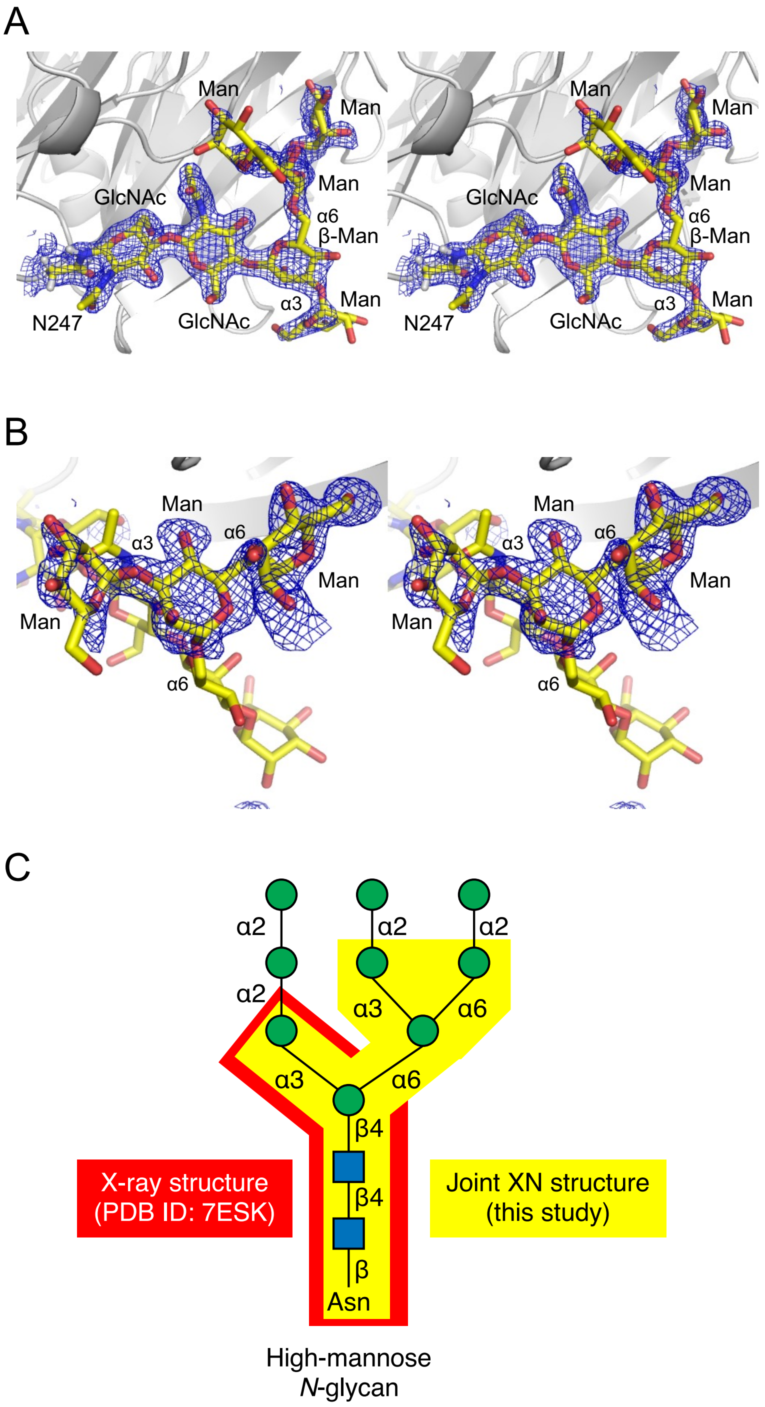


**Fig. S2.** *N*-glycan observed in the XN structure. (*A*) Stereo view of heptasaccharide (GlcNAc$_2$-Man$_5$) linked to Asn247 with $2mF_o$-$DF_c$ X-ray electron density map (1.0σ, blue mesh). (*B*) Stereo view of Man$_3$ at the α-1,6 branch with $mF_o$-$DF_c$ omit X-ray electron density map (2.7σ, blue mesh). (*C*) Schematic drawing of a typical high-mannose (oligomannose) *N*-glycan structure (1). Tetrasaccharide (GlcNAc$_2$-Man$_2$) and heptasaccharide observed in an X-ray structure and the joint XN structure are indicated, respectively. The glycan structure is represented using the Symbol Nomenclature for Glycans (SNFG) (2).

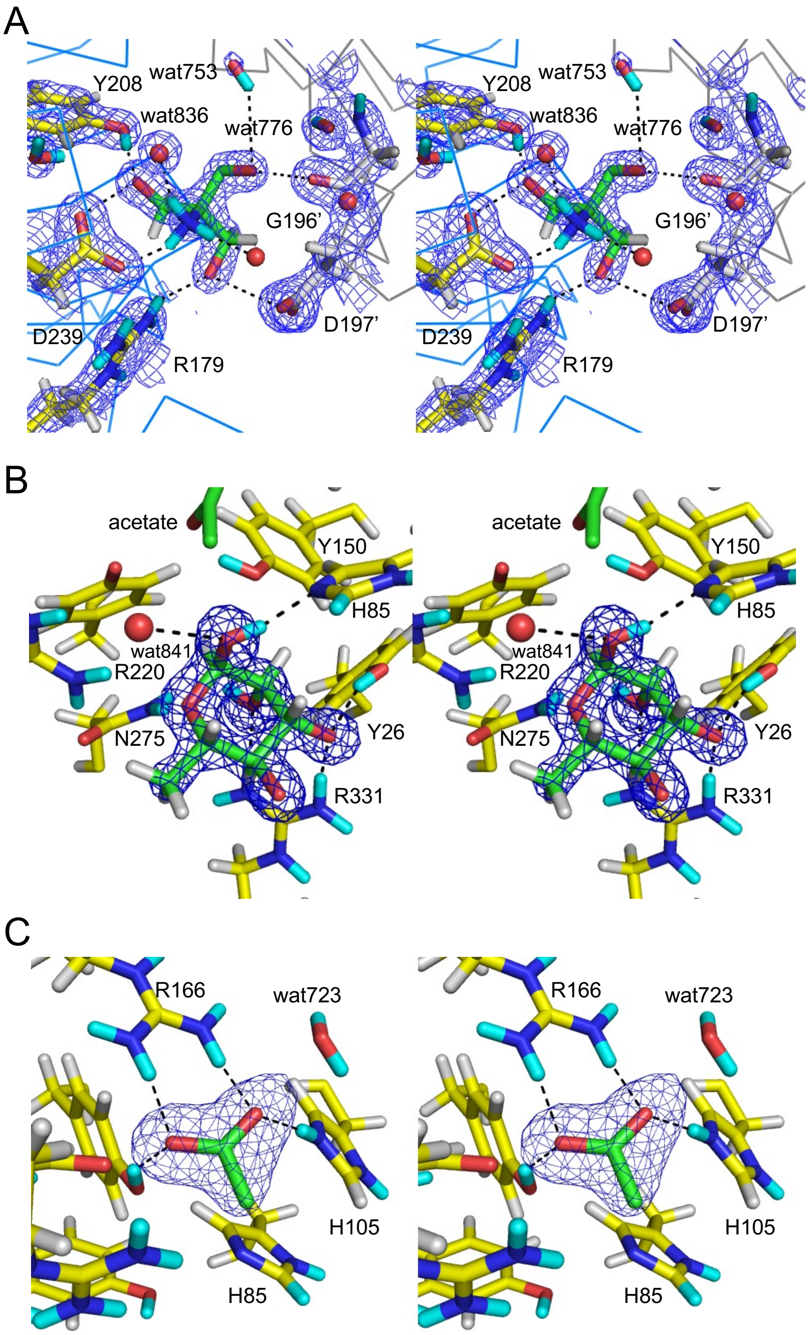


**Fig. S3.** Stereo view of ligands bound to the XN structure with X-ray electron density maps (blue mesh). H and D atoms are colored in white and cyan, respectively. Hydrogen bonds are shown with black dotted lines. (*A*) Tris-binding site with $2mF_o$-$DF_c$ map (1.5σ). Tris molecule is located at the interface of symmetry-related FoRham1 molecules. Residues from the symmetry-related molecule are indicated with prime symbols (G196’ and D197’). (*B*) Rha at subsite –1 with $mF_o$-$DF_c$ omit map (7.0σ). (C) Acetate ion at subsite +1 with $mF_o$-$DF_c$ omit map (4.0σ).

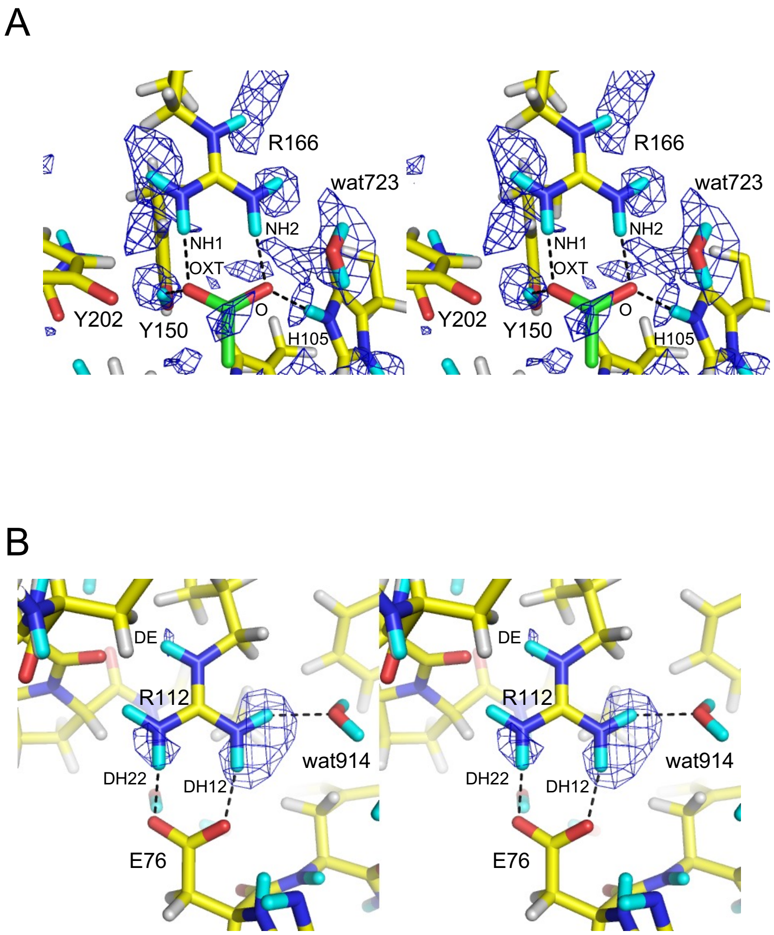


**Fig. S4.** Comparison of two Arg-carboxylate salt bridge interactions shown by stereo views with $mF_o$-$DF_c$ NSLD maps (blue mesh). H and D atoms are colored in white and cyan, respectively. Hydrogen bonds are shown with black dotted lines. (*A*) Acetate-binding site. This map is shown at a contour level of 2.4σ, which is lower than that in Fig. 2*B* (2.9σ). Following atoms were excluded from map calculation: DD1 and DE2 of His105; DH of Tyr150; DE, DH11, DH12, DH21, and DH22 of Arg166; and D1 and D2 atoms of wat52. (*B*) Arg112 and Glu76 with a map contoured at 2.8σ. Following atoms were excluded from map calculation: DE, DH11, DH12, DH21, and DH22 of Arg112.

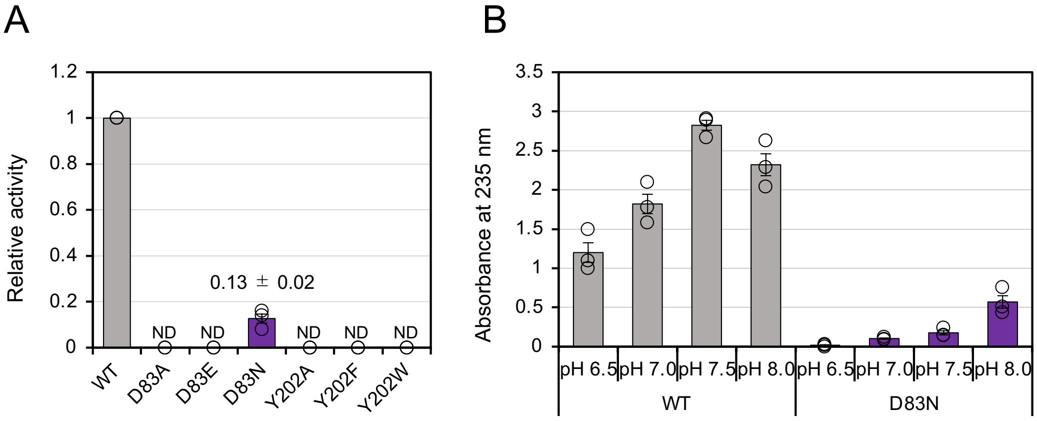


**Fig. S5.** Enzymatic activities of FoRham1 WT and mutants. (*A*) Relative activities of FoRham1 and the mutant enzymes were analyzed using GA (1%) as the substrate (n = 3). The enzymes (0.2 μM) were incubated with the substrate in 50 mM HEPES–NaOH (pH 7.0) at 30 °C for 10 min. The reaction mixture was then boiled for 3 min to inactivate the enzyme. The produced ΔGlcA was quantified at an absorbance of 235 nm. ND, not detectable. (*B*) The activities of FoRham1 and the mutant enzymes were analyzed using GA (1%) as the substrate (n = 3). The enzymes (0.2 μM) were incubated with the substrate in 50 mM MOPS–NaOH (pH 6.5–8.0) at 30 °C for 10 min. The reaction mixture was then boiled for 3 min to inactivate the enzyme. The activities were measured at an absorbance of 235 nm.

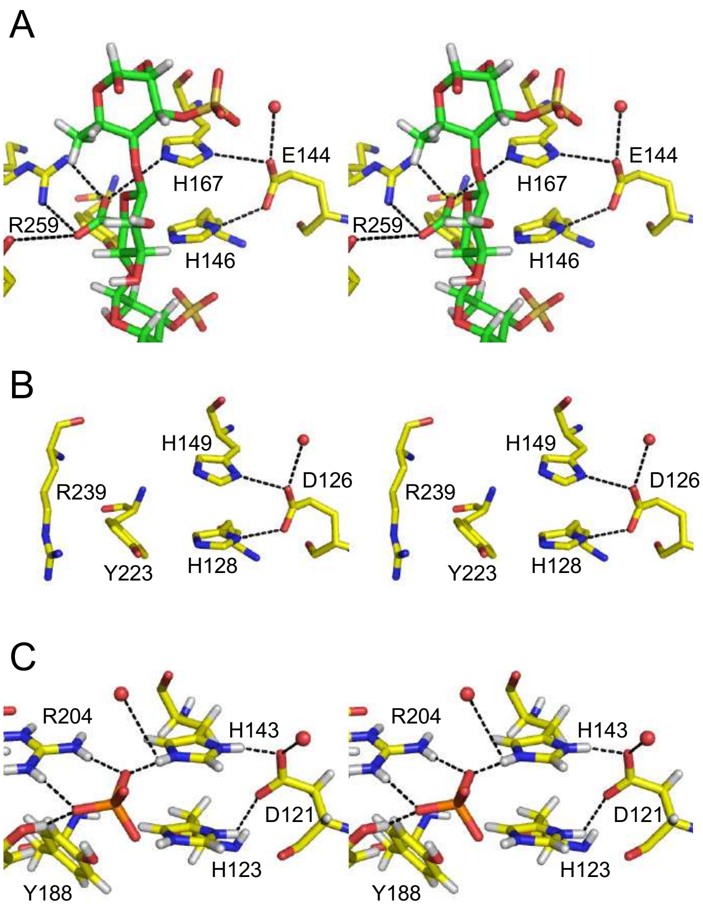


**Fig. S6.** Active sites of ulvan lyases belonging to PL24 and PL25. (*A*) PL24 LOR_107 from *Alteromonas* sp. strain LOR (PDB ID: 6BYT) complexed with a tetrasaccharide substrate (green). (*B*) PL24 Uly1 from *Catenovulum maritimum* (PDB ID: 7DRQ). (*C*) PL25 PLSV_3936 from *Pseudoalteromonas* sp. strain PLSV, (PDB ID: 5UAM) complexed with sulfate (orange). In *A* and *C*, hydrogen atoms modeled in the deposited coordinate file based on the X-ray crystal structure are shown.

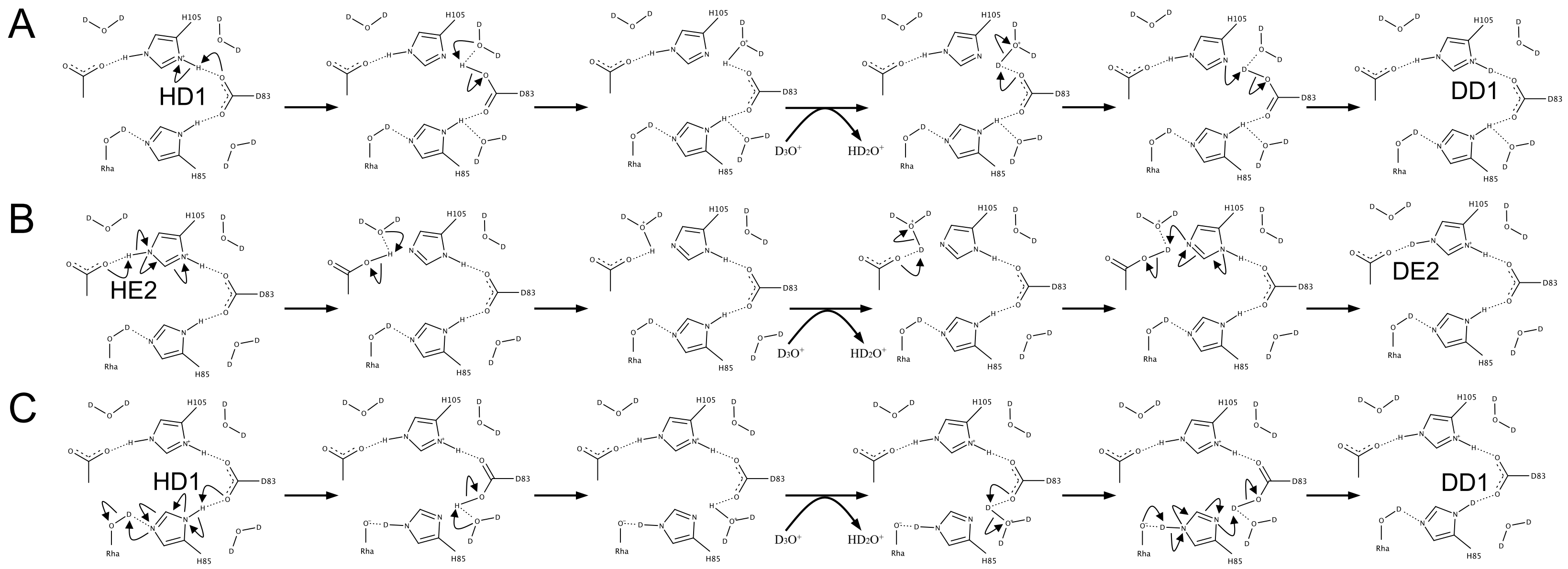

**Fig. S7.** Possible H/D exchange mechanism of (*A*) HD1 to DD1 in His105, (*B*) HE2 to DE2 in His105, and (*C*) HD1 to DD1 in His85. (*A*) HD1 of His105 initially moves to a solvent $D_2O$ molecule via OD2 of Asp83, producing $HD_2O^+$. $D3O^+$ replaces the solvent position, and a D atom moves back to ND1 of His105 via Asp83. (*B*) HE2 of His105 moves to a solvent $D_2O$ molecule via acetate oxygen, and a D atom moves back from a solvent $D3O^+$ molecule to NE2 of His105 via acetate. (*C*) Since a high-concentration stock solution of Rha dissolved in $D_2O$ was used for crystallization (see Materials and Methods), the HO1 hydrogen atom on the anomeric C1 carbon of Rha is assumed to be fully deuterated via mutarotation. HD1 of His85 moves to a solvent $D_2O$ molecule via Asp83, and a D atom moves back from the solvent $D3O^+$ molecule to ND1 of His85 via Asp83.

**Table S1.** Sequences of primers used for mutagenesis.

| Name | Sequence[a] |
|---|---|
| D83A (Fw) | 5′-ACGATGGCTGGCCATAACATGATCTCT-3′ |
| D83A (Rv) | 5′-ATGGCCAGCCATCGTCTTCTGAGTATA-3′ |
| D83E (Fw) | 5′-ACGATGGAAGGCCATAACATGATCTCT-3′ |
| D83E (Rv) | 5′-ATGGCCTTCCATCGTCTTCTGAGTATA-3′ |
| D83N (Fw) | 5′-ACGATGAACGGCCATAACATGATCTCT-3′ |
| D83N (Rv) | 5′-ATGGCCGTTCATCGTCTTCTGAGTATA-3′ |
| Y202A (Fw) | 5′-AACGCGGCTATCAACGGACTGGATTAC-3′ |
| Y202A (Rv) | 5′-GTTGATAGCCGCGTTGTTGTCGTCACC-3′ |
| Y202F (Fw) | 5′-AACGCGTTTATCAACGGACTGGATTAC-3′ |
| Y202F (Rv) | 5′-GTTGATAAACGCGTTGTTGTCGTCACC-3′ |
| Y202W (Fw) | 5′-AACGCGTGGATCAACGGACTGGATTAC-3′ |
| Y202W (Rv) | 5′-GTTGATCCACGCGTTGTTGTCGTCACC-3′ |

[a] Underlined regions indicate the mutation sites.
